# Enzyme Fragment Complementation Driven by Nucleic Acid Hybridization

**DOI:** 10.1101/2023.12.19.572427

**Authors:** Zihan Xu, Xiaoyu Zhang, Chandan Pal, Eriks Rozners, Brian P. Callahan

## Abstract

A modified protein fragment complementation assay has been designed and validated as a gain-of-signal biosensor for nucleic acid:nucleic acid interactions. The assay uses fragments of NanoBiT, the split luciferase reporter enzyme, that are esterified at their C-termini to steramers, sterol-modified oligodeoxynucleotides. The *Drosophila* hedgehog autoprocessing domain, DHhC, served as a self-cleaving catalyst for these bioconjugations. In the presence of ssDNA or RNA with segments complementary to the steramers and adjacent to one another, the two NanoBiT fragments productively associate, reconstituting NanoBiT enzyme activity. NanoBiT luminescence in samples containing nM ssDNA or RNA template exceeded background by 30-fold and as high as 120-fold depending on assay conditions. A unique feature of this detection system is the absence of a self-labeling domain in the NanoBiT bioconjugates. Eliminating that extraneous bulk broadens the detection range from short oligos to full-length mRNA.

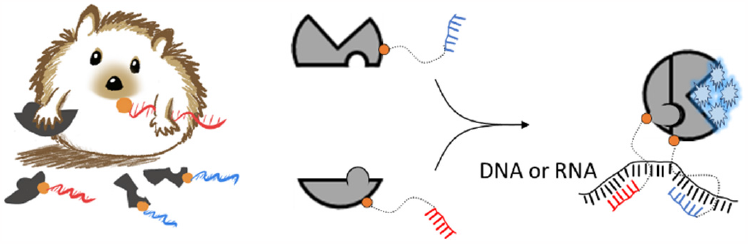

---

Proteins whose function can be turned on and off experimentally enable biosensing applications and provide valuable reagents to probe complex biological pathways. Temperature sensitive phenotypes, for example, are possible with destabilized mutant proteins that aggregate or fail to fold above some threshold temperature.[1] Domain insertion into a protein with ligand binding modules,[2] light responsive polypeptides, such as LOV2, or light responsive prosthetic groups, like azobenzene, provides an alternative and potentially reversible means for controlling protein function. [3-6]

Protein fragment complementation, the focus of the present work, is widely applied as an approach to achieve conditional activity.[7] The protein of interest is “split” into non-functional, weakly interacting fragments. Under selected conditions, those fragments are driven to assemble into a functional complex, restoring function.[8, 9] Enzymatic and nonenzymatic proteins are amenable to this type of manipulation.[10] Different triggers can be used to promote fragment complementation, such as noncovalent interactions of fused partner proteins, as in the two-hybrid assay,[11] proteolytic cleavage,[12] as well as by chemical dimerizers for concentration-responsive complementation.[13, 14]

Here we explore nucleic acid hybridization as a driver of enzyme fragment complementation with a view toward applications in RNA and DNA biosensing.[15-19] As depicted in **Scheme 1**, we envisioned a split reporter enzyme where each polypeptide fragment was covalently linked to an oligodeoxynucleotide. The sequences of those oligos would be designed for hybridization at proximal sites on a DNA or RNA of interest. In this prototype biosensor, enzyme activity reports the presence of complementary nucleic acid.

**Scheme 1.**
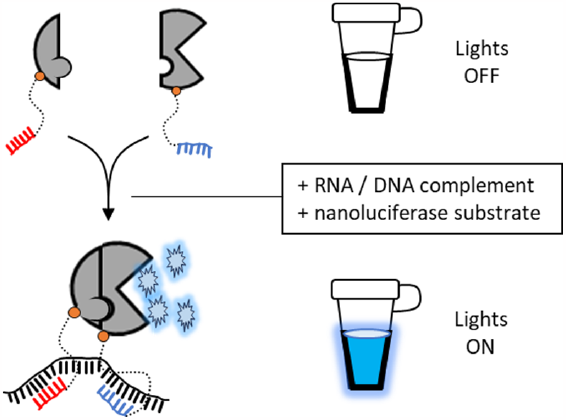
Repurposing protein fragment complementation for DNA or RNA detection. Split NanoBiT luciferase fragments with covalently attached oligonucleotides assemble via hybridization on a suitable nucleic acid template.

For evaluation, we chose the split reporter enzyme, Nano-BiT, derived by Wood and colleagues from the bioluminescent nanoluciferase. NanoBiT is comprised of a large (LrgBiT, 18 kDa) and small (SmBiT, 1.2 kDa) fragment.[20] Intrinsic affinity of LrgBiT and SmBiT is weak (K_D_ ∼ 100-200 µM), reducing spontaneous association. When LrgBiT and SmBiT are induced to associate productively at concentrations well below their K_D_ value, the two fragments reconstitute NanoBiT as an engineered heterodimer. Enzymatic oxidation of coelenterazine or its synthetic analog furimazine by NanoBiT generates glow-type luminescence. Like nanoluciferase, NanoBiT activity is independent of NTP and cofactor/coenzymes, which simplifies experimental design. Other advantages of luciferase reporters include their sensitivity, the absence of background in most biological samples and the ease of signal detection, permitting qualitative yes/no analysis even with cellphone-based imaging devices.[18, 21, 22]

To attach oligodeoxynucleotides site-specifically to LrgBiT and SmBiT with 1:1 stoichiometry we exploited the enzymatic autoprocessing domain found in *Drosophila* hedgehog protein, hereafter, DHhC. [23] The native function of DHhC is to cleave off and covalently attach cholesterol to an adjacent N-terminal polypeptide within a precursor form of hedgehog. [24, 25] We and others have found that DHhC is substrate promiscuous, retaining cleavage/sterylation activity when fused to heterologous N-terminal polypeptides (e.g., nanoluciferase, maltose binding protein, lysozyme, cyan fluorescent protein, etc.) and accepting a broad range of cholesterol analogs as alternative substrates, including monosterylated oligonucleotides, or steramers.[23, 25, 26] In a steramer, the sterol’s fused ring structure is maintained for substrate recognition by DHhC while the iso-octyl sterol side chain is modified to serve as a linker for covalent oligo attachment (*below*). DHhC is active toward steramer substrates with oligos of varying sequence, length, chemical modification and secondary structure.[23] We recently reported a cost-effective means for preparing steramers by solid phase synthesis using commercial reagents.[27]

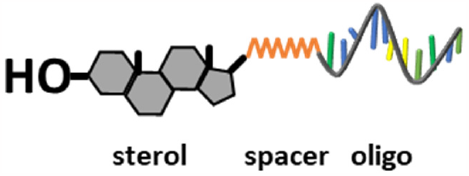

The general method for protein-to-steramer bioconjugation using DHhC resembles other self-labeling proteins such as HaloTag or HUH tag.[28, 29] The key distinction is that DHhC self-cleaves from the product simultaneously with the coupling event whereas self-labeling domains remain attached. A chimeric gene construct is assembled first in which the protein of interest (POI) is fused at its C-terminus to DHhC. A single glycine residue represents the minimal spacer between the protein of interest and the first residue of DHhC. The alpha carboxyl group of that glycine spacer residue will be the site of sterol esterification. Following expression of the chimeric POI-DHhC gene under sterol-free conditions, such as in *E. coli*, the protein precursor is activated by addition of steramer. No ATP or cofactors are involved. As mentioned above, protein-steramer bioconjugation displaces DHhC, furnishing an almost scar-free product (**Figure 1**). Note that, here, we incorporated (Gly-Ser) C-terminal spacer to allow for flexibility in split NanoBiT re-assembly. [20, 30, 31]

**Figure 1.**
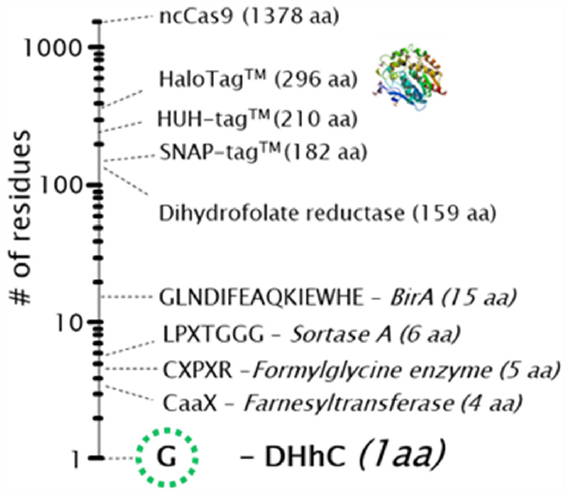
Extraneous peptide sequences or self-labeling domains that remain on the selected bioconjugate product cover a wide range. *Drosophila* Hedgehog HhC (DHhC) catalyzed protein-nucleic acid conjugation, used in the present study, leaves a small “scar” of a single glycine residue.

The chemi-enzymatic approach to generate the two NanoBiT steramer components of this prototype nucleic acid biosensor is summarized in **Figure 2**. For the NanoBiT components, we designed constructs where LrgBiT and SmBiT were fused to DHhC, an N-terminal SUMO tag was incorporated as a solubility enhancer[32] and a C-terminal His_6_ tag was added to DHhC for precursor protein purification. The two constructs, SUMO-LrgBiT-DHhC-His_6_ and SUMO-SmBiT-DHhC-His_6_, were overexpressed in *E. coli* in soluble form and purified under native conditions by Ni-NTA chromatography (**Supporting Figure 1**).

**Figure 2.**
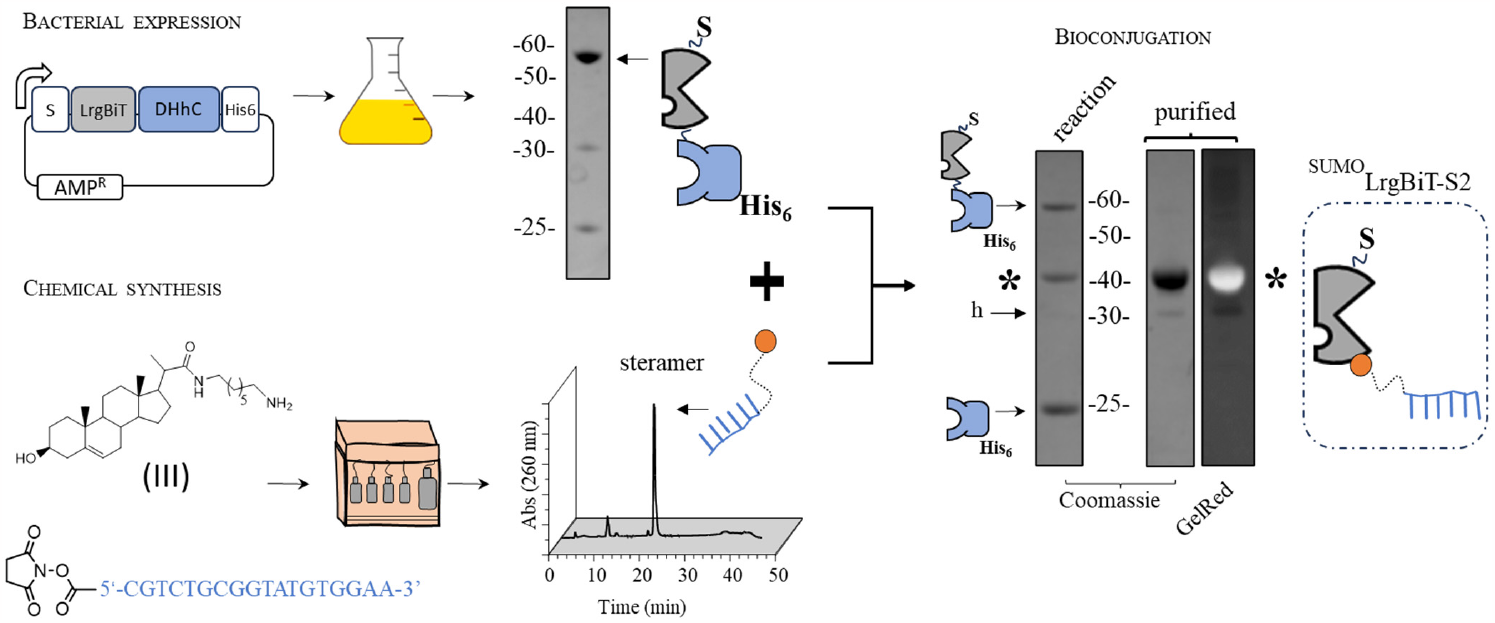
Convergent chemi-enzymatic synthesis of NanoBiT fragment-oligonucleotide conjugates. (Top) Design, *E. coli* expression and purification of LrgBiT fusion with the bioconjugating domain, DHhC. Constructs contain an N-terminal SUMO (S) protein to enhance solubility and a C-terminal hexahistidine tag (His_6_) for Ni-NTA chromatography. (Bottom) Steramer preparation. Monosterylation of oligodeoxynucleotides was carried out by amide coupling between (III) and NHS-ester modified oligodeoxynucleotides, followed by purification with reverse phase chromatography. (Side right) Bioconjugation. Precursor protein, ^SUMO^LrgBiT-DHhC-His_6_ is combined with steramer, leading to the displacement of DHhC-His_6_ and covalent steramer attachment to ^SUMO^LrgBiT, to yield ^SUMO^LrgBiT-S2 (*). Competing side reaction where water replaces the steramer in the bioconjugation is minimal (h, product from hydrolysis). Bioconjugates were purified by agarose gel extraction, then concentrated by spin column.

For proof of concept, oligo sequences for two steramers (**Table 1**) were designed for annealing at adjacent sites within the NSP10 gene from SARS-CoV-2. NSP10 encodes a protein cofactor of the viral 2’-O-RNA methyltransferase.[33, 34] Unlike variability reported in other SARS-CoV-2 coding sequences, NSP10 appears relatively stable,[35, 36] making this region an attractive target for molecular detection. Steramers were prepared in two steps, first joining heptanediamine to 23,24-bisnor-5-cholenic acid-3β-ol to produce **III**, followed by amide coupling of **III** to resin-bound oligodeoxynucleotide equipped with 5’ NHS ester modification. After deprotection and resin cleavage, the two steramers, S1 and S2, were desalted and isolated by reverse phase chromatography.

**Table 1.**
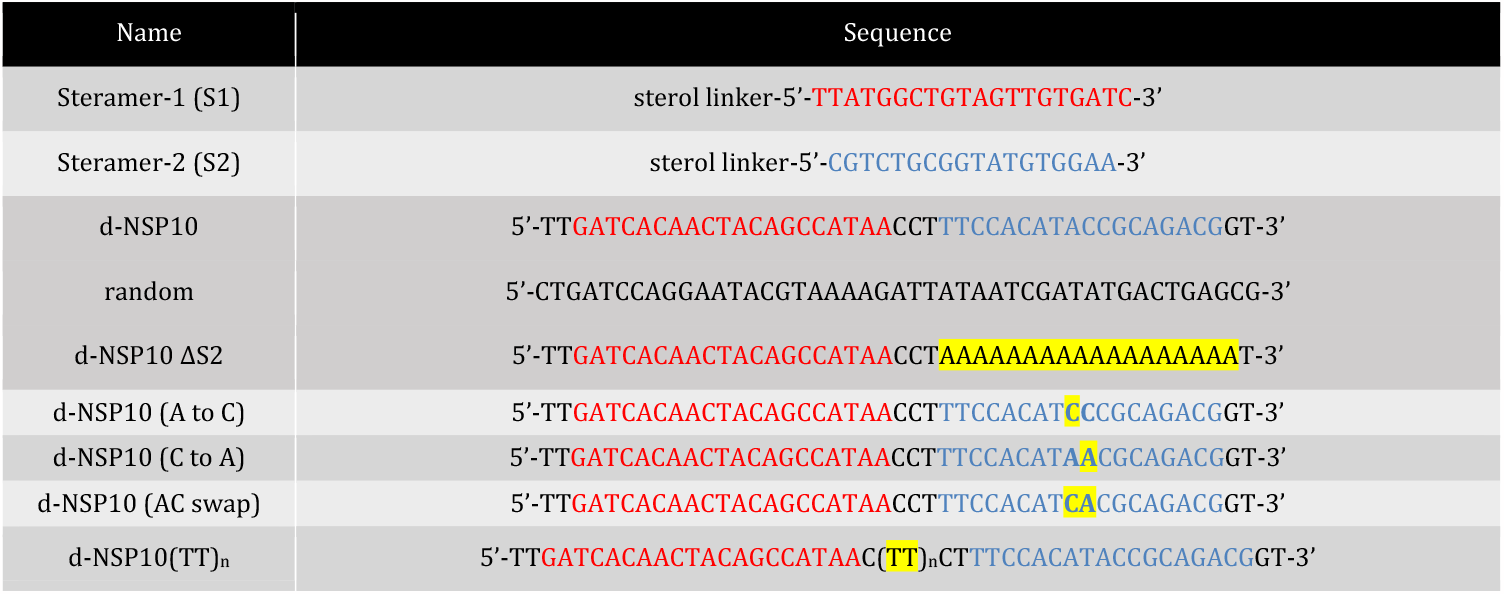
Sequences of steramer 1 and 2 and nucleic acid targets evaluated here for potential hybridization-driven protein fragment complementation.

Covalent steramer attachment by DHhC to the NanoBiT fragments was monitored by denaturing SDS-PAGE followed by staining with Gel-Red (nucleic acid specific) and Coomassie blue. In a typical experiment, reactions were initiated on the benchtop at 23 °C in Bis-Tris buffered solution (pH 7.1) containing 1.5 µM precursor protein and 50 µM steramer. Initial steramer concentration represents 2x the K_M_ value with DHhC, determined separately using a FRET-based DHhC activity assay (**Supporting Figure 2**).[26] Analysis of the reaction mixture by denaturing SDS-PAGE showed the consumption of precursor protein along with the accumulation of the respective bioconjugate. **Figure 2**, right side, is a representative gel for the SUMO-LrgBiT coupling to S2. The calculated molecular weight is 36.6 kDa for the product, which agrees well with the extrapolated value of 37.1 kDa, based on comparison to the MW ladder. This species is visible with Coomassie staining and the nucleic acid stain Gel-Red. Only trace amounts of the hydrolytic un-conjugated side-product, SUMO-LrgBiT, was observed (**Figure 2**, see gel, “h”). Reactions ranged from 50-80% conversion, with highest yields coming from precursor protein that was isolated by Ni-NTA chromatography then further purified by SEC. Agarose gel extraction offered a convenient and inexpensive means for the final isolation step (**Supporting Figure 3**).

Robust NanoBit luminescence signal was observed when the two bioconjugates, hereafter ^SUMO^Sm-S1 and ^SUMO^Lg-S2, were combined with a complementary single stranded NSP10 oligodeoxynucleotide, d-NSP10 (**Figure 3A**). NanoBiT complementation was assayed in solutions containing ^SUMO^Sm-S1 and ^SUMO^Lg-S2 and d-NSP10 added together in a ratio of 1:1:1 over a dilution series from 1 × 10^−7^ M to 48 × 10^−12^ M. As a control, we measured spontaneous NanoBiT complementation from ^SUMO^Sm-S1 and ^SUMO^Lg-S2 over that same concentration range in the absence of d-NSP10 template (**Supporting Figure 4**). Additional negative control experiments were carried out using ^SUMO^Sm-S1 and ^SUMO^Lg-S2 mixed with ssDNA template of the same length as d-NSP10 but randomized sequence; samples with the random oligo generated NanoBiT signal equal to the “no template” readings (**Figure 3A**, “random”). We also attempted to measure NanoBiT reconstitution with a ssDNA template where the hybridization site for ^SUMO^Lg-S2 was mutated to a poly T track; luminescence again matched the background (no template) readings (**Figure 3A**, “d-NSP10 ΔS2”). Based on those results, we selected 25 × 10^−9^ M as the working concentrations of ^SUMO^Sm-S1 and ^SUMO^Lg-S2. This setup, which is similar to conditions used in related NanoBiT complementation assays (**Supporting Table 1**), conserved NanoBiT-steramer components while providing maximum signal / background of 30-60 fold and a lower limit for complementary template detection of 2 ×10^−9^ M.

**Figure 2.**
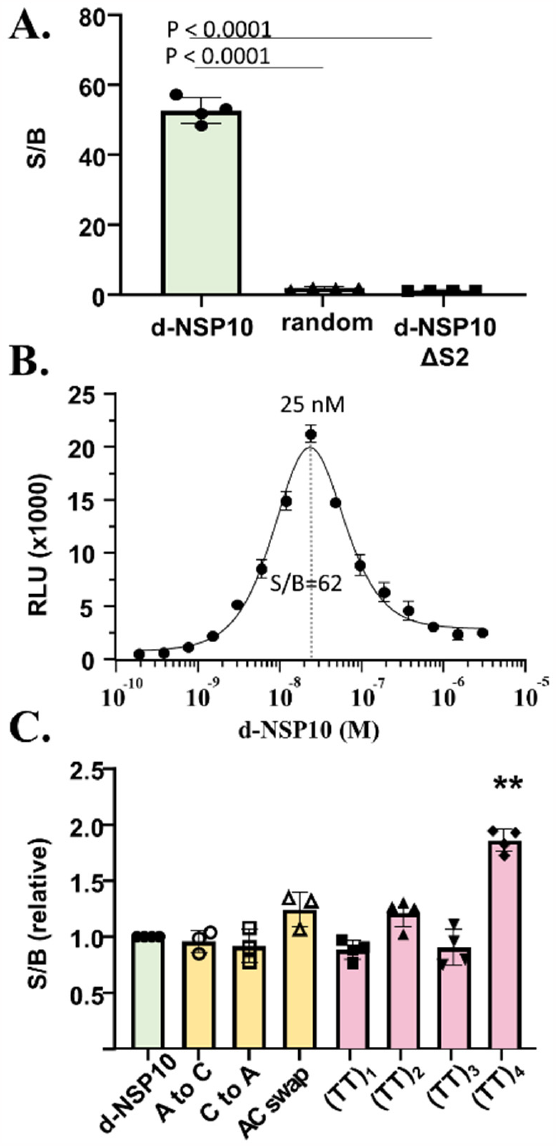
NanoBiT fragment-steramer bioconjugates enable nucleic acid hybridization driven protein fragment complementation. A. NanoBiT signal generated from samples containing the indicated ssDNA templates. Oligonucleotide templates, ^SUMO^Sm-S1, ^SUMO^Lg-S2 were (1:1:1), 25 × 10^−9^ M, final. Statistical analysis: one-way ANOVA followed by T-test (GraphPad). **B**. Hook-effect in concentration-response plot with increasing d-NSP10. **C**. Assessing NanoBiT luminescence with template variants. Conditions same as (A). Only (TT)_4_ showed a significant difference from control, d-NSP10. **, P<.0001

As shown in **Figure 3B**, luminescence from reconstituted NanoBiT plotted as a function of increasing d-NSP10 template yielded a bell-shaped curve. “Hook-effect” behavior is consistent with non-cooperative tripartite interactions where one component of the assembly provides a scaffold to bring the remaining two components together.[18, 37]Here, d-NSP10 is the scaffolding element. Signal/background was highest when the three components were equimolar. To the right of the equivalence point, the excess template favors formation of enzymatically in-active binary complexes: d-NSP10 hybridized with ^SUMO^Sm-S1 only or with ^SUMO^Lg-S2 only.

We found that NanoBiT reconstitution remained durable with several “mutant” d-NSP10 templates (**Figure 3C**). We first tested templates where the ^SUMO^Lg-S2 site contained single or double nucleotide substitutions (see **Table 1**: d-NSP10 (A to C); d-NSP10 (C to A); d-NSP10 (AC swap)). We observed only modest changes in luminescence, < 1.5-fold, compared to the positive control, d-NSP10, **Figure 3C** (yellow). Tolerance toward base-pair mismatches is not unexpected for a linear hybridization involving oligos of this length.[38, 39] In a related split enzyme system, using oligodeoxynucleotide-modified fragments of dihydrofolate reductase, split enzyme reassembly could be driven by hybridization to a ssDNA template carrying as many as five mismatches (**Supporting Table 1**).[19]

Next, we tested templates in which the hybridization sites of ^SUMO^Sm-S1 and ^SUMO^Lg-S2 were perfectly complementary but spread farther apart by insertions of 2, 4, 6, or 8 deoxy thymidine nucleotides (**Table 1**. See d-NSP10(TT)_n_). In samples containing the spacer templates d-NSP10(TT)_1,_ d-NSP10(TT)_2_ and d-NSP10(TT)_3_ we observed NanoBiT signal indistinguishable from samples mixed with d-NSP10 **Figure 3C** (pink). Only d-NSP10(TT)_4_ stood out in this comparison, and interestingly, the luminescence was enhanced, suggesting that there is room for further design optimization. In summary, these results indicate that there is tolerance toward single and double nucleotide substitutions in the nucleic acid target sequence and that there is flexibility in the distance separating the sites for hybridization.

With the modified NanoBiT complementation system working largely as intended on ssDNA, we turned our attention to the detection of target RNA. A synthetic SARS-CoV-2 NSP10 gene (460 bp) was cloned into the pGEM-3 vector and transcribed in vitro in the forward and reverse directions to generate sense (+) and anti-sense (-) NSP10 RNA transcripts. Samples containing ^SUMO^Sm-S1, ^SUMO^Lg-S2 and equivalent concentration (25 × 10^−9^ M) of the purified antisense transcript generated strong NanoBiT signal (**Figure 4**). RLU values were on par with those containing d-NSP10, used a positive control. A hook-effect in concentration-response plots was also apparent (**Supporting Figure 5**). By contrast, NanoBiT reconstitution was not supported by samples containing the NSP10 sense transcript, which harbors segments identical to the oligonucleotides in ^SUMO^Sm-S1 and ^SUMO^Lg-S2. In summary, the split NanoBiT-steramer conjugates provide hybridization-driven luminescence detection of ssDNA and RNA.

**Figure 3.**
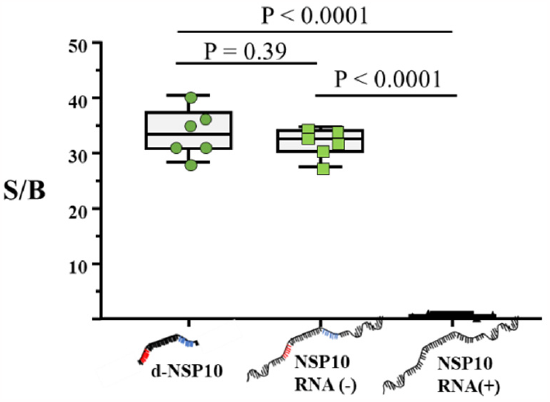
NanoBiT reconstitution reports the presence of SARS-CoV-2 NSP-10 RNA. In vitro transcribed anti-sense NSP10 RNA (squares) but not NSP10 sense RNA (triangles) triggers NanoBiT activity. Synthetic d-NSP10 was added as a positive control (circles). Nucleic acid and NanoBiT components were added (1:1:1) to 25 ×10^−9^ M, final. Statistical analysis: one-way ANOVA followed by T-test (GraphPad).

As this work was underway, a conceptually similar nucleic acid biosensor was reported that involves fusions of the NanoBiT fragments to a noncatalytic Cas9 protein (dCas9).[18, 40] In that system, the ∼160 kDa dCas9 provided the linker to (non-covalently) join LrgBiT and SmBiT to an RNA oligonucleotide in a manner similar to the way the sterol molecule here covalently links ^SUMO^LrgBiT and ^SUMO^SmBiT to DNA oligonucleotides. Although the assay also suffered from a hook-effect, a practical advantage of using dCas9 is that the guide RNAs for dsDNA template recognition do not require chemical modification. One potential drawback is the restricted hybridization specificity of this large ribonucleoprotein complex: split NanoBiT reconstitution required that the target harbor a 5′-NGG-3′ PAM and its reverse complement 5′-CCN-3′ within 27–52 nt of each other. The shortest dsDNA template that supported NanoBiT reconstitution with the dCas9 linker was 450 bp. Nonetheless, once optimized, the dCas9/NanoBiT “LUNAS” method from Merkx and colleagues[18] showed impressive sensitivity and appeared amenable to translation as a point-of-care diagnostic. Specificity of LUNAS to discriminate nucleotide mismatches was not disclosed.

In this study, we designed, prepared and validated a prototype split enzyme complementation assay driven by nucleic acid hybridization. We attached monosterylated oligodeoxynucleotides (steramers) site-specifically to the carboxy termini of SUMO-fused LrgBiT and SmBiT fragments using the self-cleaving *Drosophila* hedgehog autoprocessing domain, DHhC. It seems reasonable to assume that direct covalent attachment of oligonucleotide to split enzyme fragments by a small molecule linker, rather than through a self-labeling macromolecule, would moderate steric interference to productive protein fragment complementation. Moreover, small molecule linkers may be beneficial for split protein assembly on shorter nucleic acids of interest including non-coding RNA. Although the solid phase synthesis of steramers could present a practical obstacle to the application of DHhC bioconjugation, an alternative solution phase steramer preparation has been reported involving straight-forward oxime chemistry.[23]

Next generation nucleic acid biosensor assays of the general type described here will need to address limitations in mismatch discrimination as well as the hook-effect in concentration response plots, the latter causing diminished signal output with super stoichiometric template. We anticipate that using hairpin oligo components in place of the linear oligos, used here, will increase the sequence specificity of protein fragment complementation, as it does with molecular beacons.[41, 42] Protein-steramer bioconjugation by DHhC proceeds unimpeded with hairpin oligos, making this an attractive test system.[27] The second challenge posed by the hook-effect is a design defect that threatens assay reliability through false negatives, where increasing nucleic acid template results in decreasing protein fragment complementation. Engineering cooperativity into the NanoBiT/template assembly offers one solution.[43] A split enzyme-steramer complementation assay with cooperative 1:1:1 template hybridization would drive reporter enzyme reconstitution even with a template surplus.

## ASSOCIATED CONTENT

### Supporting Information

vector construction, protein expression, steramer kinetic assays, chemical synthesis, NMR and MALDI-TOF. This material is available free of charge via the Internet at http://pubs.acs.org.

## AUTHOR INFORMATION

### Author Contributions

The manuscript was written through contributions of all authors. All authors have given approval to the final version of the manuscript.

### Funding Sources

We acknowledge generous support from the National Institute of Allergy and Infectious Diseases (Grant R03AI163907 to B.P.C.), and the National Institute of General Medical Sciences (Grant R35GM130207 to E.R.).

## ABBREVIATIONS

DHhC: Drosophila hedgehog C-terminal domain
SUMO: small ubiquitin modifier protein
MW: molecular weight
ssDNA: single stranded DNA
dsDNA: double stranded DNA
NSP10: non-structural protein 10
Ni-NTA: Nickel (2+)-Nitriloacetic acid
S1: steramer 1
S2: steramer 2
NSP: nonstructural protein

## Notes

### Competing Interest Statement

The authors have declared no competing interest.

## REFERENCES

1. Ma, C., et al., The segment polarity gene hedgehog is required for progression of the morphogenetic furrow in the developing Drosophila eye. Cell, 1993. 75(5): p. 927–38.

2. Skretas, G. and D.W. Wood, Regulation of protein activity with small-molecule-controlled inteins. Protein Sci, 2005. 14(2): p. 523–32.

3. Dagliyan, O., et al., Engineering extrinsic disorder to control protein activity in living cells. Science, 2016. 354(6318): p. 1441–1444.

4. Blacklock, K.M., et al., Computational Design of a Photocontrolled Cytosine Deaminase. J Am Chem Soc, 2018. 140(1): p. 14–17.

5. Szymanski, W., et al., Reversible photocontrol of biological systems by the incorporation of molecular photoswitches. Chem Rev, 2013. 113(8): p. 6114–78.

6. Wu, Y.I., et al., Spatiotemporal control of small GTPases with light using the LOV domain. Methods Enzymol, 2011. 497: p. 393–407.

7. Michnick, S.W., Exploring protein interactions by interaction-induced folding of proteins from complementary peptide fragments. Curr Opin Struct Biol, 2001. 11(4): p. 472–7.

8. Johnsson, N. and A. Varshavsky, Split ubiquitin as a sensor of protein interactions in vivo. Proc Natl Acad Sci U S A, 1994. 91(22): p. 10340–4.

9. Rossi, F., C.A. Charlton, and H.M. Blau, Monitoring protein-protein interactions in intact eukaryotic cells by beta-galactosidase complementation. Proc Natl Acad Sci U S A, 1997. 94(16): p. 8405–10.

10. Banaszynski, L.A. and T.J. Wandless, Conditional control of protein function. Chem Biol, 2006. 13(1): p. 11–21.

11. Fields, S. and O. Song, A novel genetic system to detect protein-protein interactions. Nature, 1989. 340(6230): p. 245–6.

12. Callahan, B.P., M.J. Stanger, and M. Belfort, Protease activation of split green fluorescent protein. Chembiochem, 2010. 11(16): p. 2259–63.

13. Remy, I. and S.W. Michnick, Clonal selection and in vivo quantitation of protein interactions with protein-fragment complementation assays. Proc Natl Acad Sci U S A, 1999. 96(10): p. 5394–9.

14. Mootz, H.D., et al., Conditional protein splicing: a new tool to control protein structure and function in vitro and in vivo. J Am Chem Soc, 2003. 125(35): p. 10561–9.

15. Demidov, V.V., et al., Fast complementation of split fluorescent protein triggered by DNA hybridization. Proc Natl Acad Sci U S A, 2006. 103(7): p. 2052–6.

16. Furman, J.L., et al., Toward a general approach for RNA-templated hierarchical assembly of split-proteins. J Am Chem Soc, 2010. 132(33): p. 11692–701.

17. Zhou, L., et al., Tandem reassembly of split luciferase-DNA chimeras for bioluminescent detection of attomolar circulating microRNAs using a smartphone. Biosens Bioelectron, 2021. 173: p. 112824.

18. van der Veer, H.J., et al., Glow-in-the-Dark Infectious Disease Diagnostics Using CRISPR-Cas9-Based Split Luciferase Complementation. ACS Cent Sci, 2023. 9(4): p. 657–667.

19. Oltra, N.S., J. Bos, and G. Roelfes, Control over enzymatic activity by DNA-directed split enzyme reassembly. Chembiochem, 2010. 11(16): p. 2255–8.

20. Dixon, A.S., et al., NanoLuc Complementation Reporter Optimized for Accurate Measurement of Protein Interactions in Cells. ACS Chem Biol, 2016. 11(2): p. 400–8.

21. Sekhon, H. and S.N. Loh, Engineering protein activity into off-the-shelf DNA devices. Cell Reports Methods, 2022. 2(4).

22. Chang, D., et al., Smartphone DNA or RNA Sensing Using Semisynthetic Luciferase-Based Logic Device. ACS Sens, 2020. 5(3): p. 807–813.

23. Zhang, X., et al., Protein-Nucleic Acid Conjugation with Sterol Linkers Using Hedgehog Autoprocessing. Bioconjug Chem, 2019. 30(11): p. 2799–2804.

24. Porter, J.A., K.E. Young, and P.A. Beachy, Cholesterol modification of hedgehog signaling proteins in animal development. Science, 1996. 274(5285): p. 255–9.

25. Hall, T.M., et al., Crystal structure of a Hedgehog autoprocessing domain: homology between Hedgehog and self-splicing proteins. Cell, 1997. 91(1): p. 85–97.

26. Owen, T.S., et al., Forster resonance energy transfer-based cholesterolysis assay identifies a novel hedgehog inhibitor. Anal Biochem, 2015. 488: p. 1–5.

27. Zhang, X., et al., Enzymatic Beacons for Specific Sensing of Dilute Nucleic Acid. Chembiochem, 2022. 23(4): p. e202100594.

28. Los, G.V., et al., HaloTag: a novel protein labeling technology for cell imaging and protein analysis. ACS Chem Biol, 2008. 3(6): p. 373–82.

29. Lovendahl, K.N., A.N. Hayward, and W.R. Gordon, Sequence-Directed Covalent Protein-DNA Linkages in a Single Step Using HUH-Tags. J Am Chem Soc, 2017. 139(20): p. 7030–7035.

30. Cooley, R., et al., Development of a cell-free split-luciferase biochemical assay as a tool for screening for inhibitors of challenging protein-protein interaction targets. Wellcome Open Res, 2020. 5: p. 20.

31. Chretien, A.E., et al., Extended Linkers Improve the Detection of Protein-protein Interactions (PPIs) by Dihydrofolate Reductase Protein-fragment Complementation Assay (DHFR PCA) in Living Cells. Mol Cell Proteomics, 2018. 17(2): p. 373–383.

32. Butt, T.R., et al., Ubiquitin fusion augments the yield of cloned gene products in Escherichia coli. Proc Natl Acad Sci U S A, 1989. 86(8): p. 2540–4.

33. Lin, S., et al., Crystal structure of SARS-CoV-2 nsp10/nsp16 2’-O-methylase and its implication on antiviral drug design. Signal Transduct Target Ther, 2020. 5(1): p. 131.

34. Krafcikova, P., et al., Structural analysis of the SARS-CoV-2 methyltransferase complex involved in RNA cap creation bound to sinefungin. Nat Commun, 2020. 11(1): p. 3717.

35. Zhou, L., et al., Programmable low-cost DNA-based platform for viral RNA detection. Sci Adv, 2020. 6(39).

36. Tasakis, R.N., et al., SARS-CoV-2 variant evolution in the United States: High accumulation of viral mutations over time likely through serial Founder Events and mutational bursts. PLoS One, 2021. 16(7): p. e0255169.

37. Douglass, E.F., Jr., et al., A comprehensive mathematical model for three-body binding equilibria. J Am Chem Soc, 2013. 135(16): p. 6092–9.

38. Aboul-ela, F., et al., Base-base mismatches. Thermodynamics of double helix formation for dCA3XA3G + dCT3YT3G (X, Y = A,C,G,T). Nucleic Acids Res, 1985. 13(13): p. 4811–24.

39. Urakawa, H., et al., Optimization of single-base-pair mismatch discrimination in oligonucleotide microarrays. Appl Environ Microbiol, 2003. 69(5): p. 2848–56.

40. Liu, L., Y. Wang, and D.L. Schmitt, LUNAS: A Rapid and Sensitive Nucleic Acid Detection Assay Using Split NanoLuc Luciferase Complementation. ACS Cent Sci, 2023. 9(4): p. 593–596.

41. Tyagi, S. and F.R. Kramer, Molecular beacons: probes that fluoresce upon hybridization. Nat Biotechnol, 1996. 14(3): p. 303–8.

42. Hunt, E.A. and S.K. Deo, Bioluminescent stem-loop probes for highly sensitive nucleic acid detection. Chem Commun (Camb), 2011. 47(33): p. 9393–5.

43. Liu, S., et al., Rational Screening for Cooperativity in Small-Molecule Inducers of Protein-Protein Associations. J Am Chem Soc, 2023. 145(42): p. 23281–23291.

