## Supplementary Information for "Enzyme Fragment Complementation Driven by Nucleic Acid Hybridization"

### SUPPORTING FIGURES

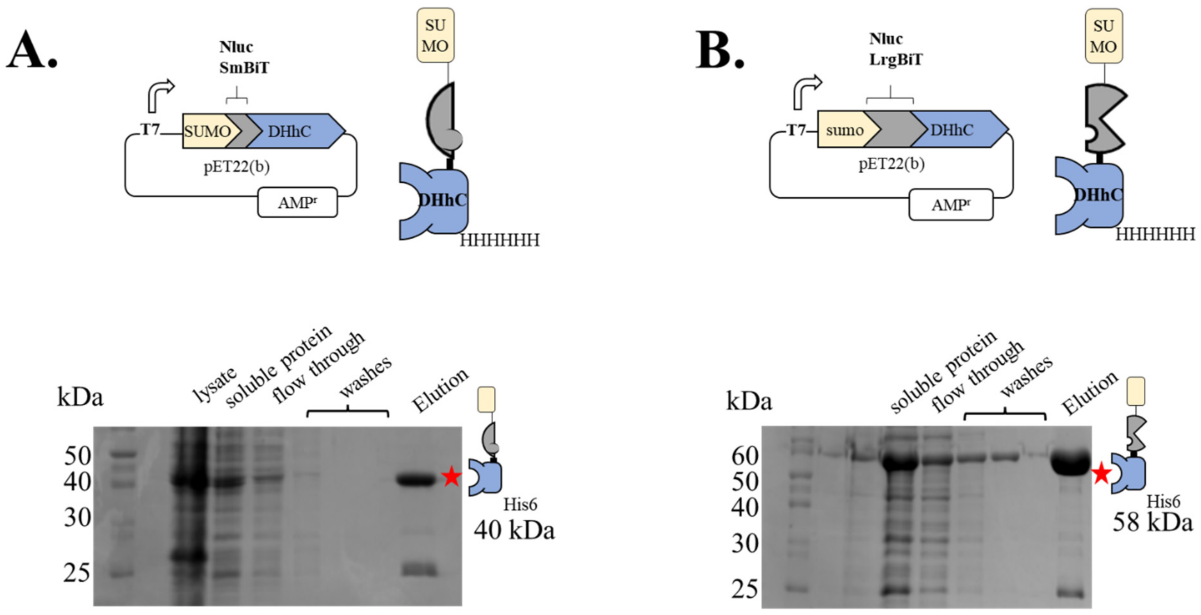

**Supporting Figure 1.** Bioconjugation fusion constructs of LrgBiT and SmBiT are over expressed in *E. coli* in soluble form and purified by Ni-NTA chromatography. (A) SUMO-SmBiT-DHhC-His$_6$ expression vector and representative analysis of *E. coli* expression and Ni-NTA purification by denaturing SDS-PAGE. (B) SUMO-LrgBiT-DHhC-His$_6$ expression vector and representative analysis of *E. coli* expression / Ni-NTA purification by denaturing SDS-PAGE.

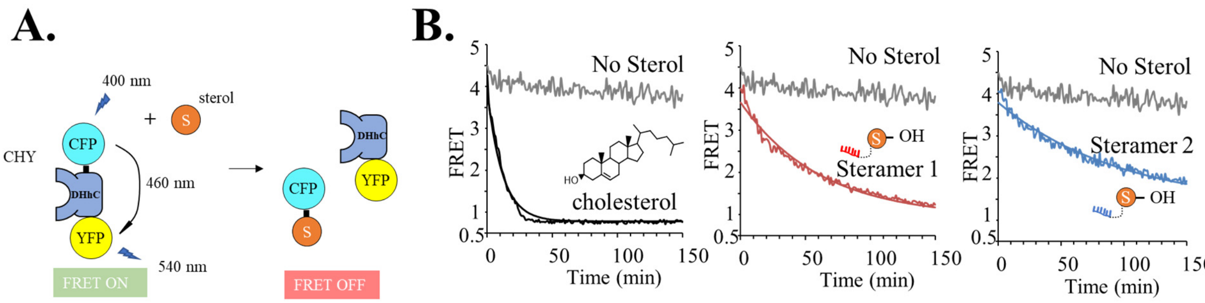

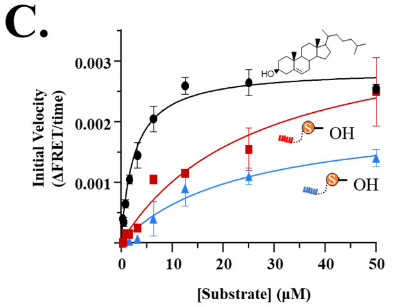

D.

| 50 μM | cholesterol | Steramer 1 | Steramer 2 |
|---|---|---|---|
| $k_{obs}$ ($s^{-1}$x $10^{-3}$) | 1.5 | 0.27 | 0.17 |
| $K_M$ (μM) | 2.57 | 26.3 | 23.7 |

**Supporting Figure 2.** Steramer 1 and 2 are alternative substrates for DHhC. (A) FRET-reporter protein, C-H-Y, to monitor DHhC catalyzed sterolysis in real time. (B) Kinetic traces with C-H-Y using the native substrate, cholesterol (positive control), alongside kinetic traces with steramer 1 and steramer 2. (C) Michaelis Menten plots of reaction velocity as a function of increasing substrate concentration. Rates calculated using the FRET reporter. Cholesterol included as a comparator. (D) Summary of kinetic constants for substrate cholesterol, steramer 1 and steramer 2.

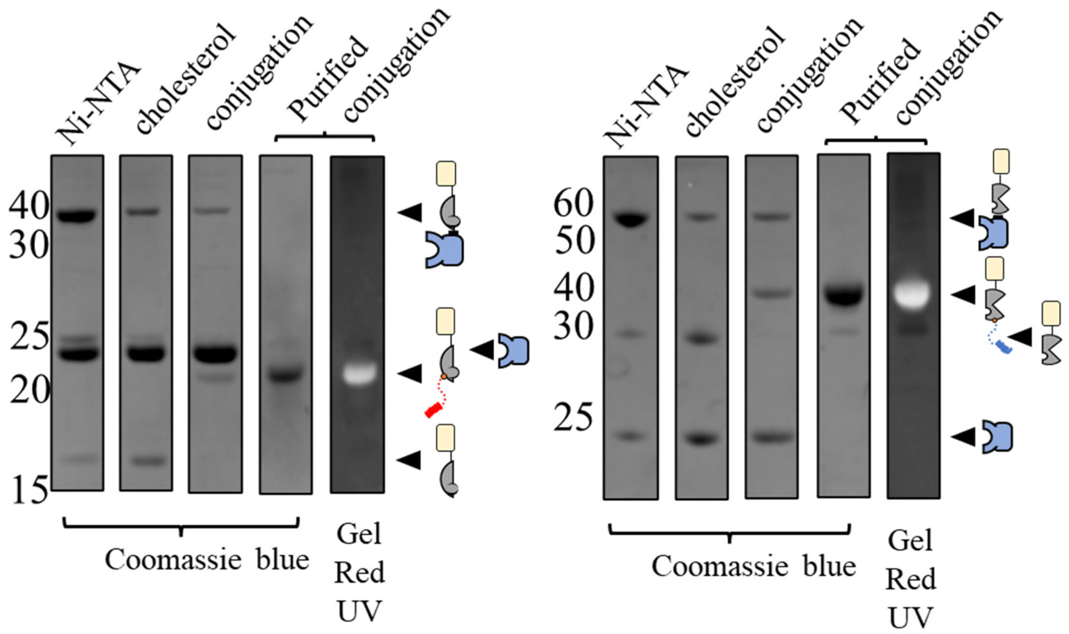

**Supporting Figure 3.** Steramer bioconjugation of SUMO-SmBiT (A) and SUMO-LrgBiT (B) catalyzed by DHhC. Reaction progress and sample purity judged by denaturing SDS-PAGE (12%). For each data set, the lane order is as follows: Lane 1, NiNTA purified precursor protein; Lane 2, DHhC catalyzed reaction with native substrate, cholesterol (positive control); Lane 3. DHhC catalyzed bioconjugation with steramer; Lane 4. Agarose purified, concentrated bioconjugate, SUMO-(NanoBiT fragment)-steramer Lane 5. Same as lane 4 except visualized by GelRed stain for nucleic acid.

**A.**

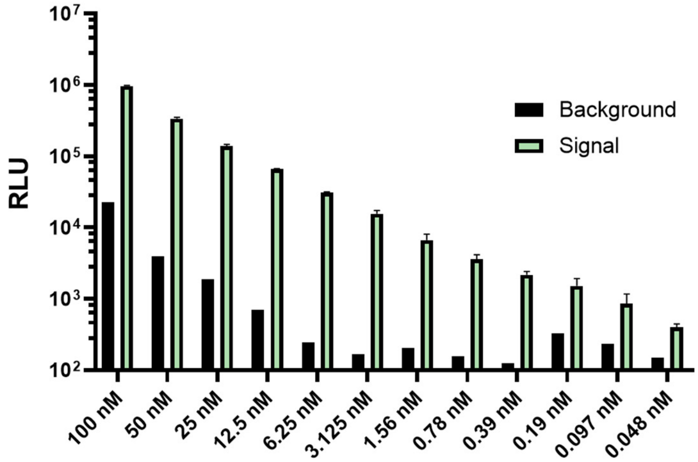

**B.**

| | BACKGROUND RLU | SIGNAL RLU | SIGANL/BACKGROUND RLU |
|---|---|---|---|
| 100 nM | 22441 | 959700 | 43 |
| 50 nM | 3930 | 336773 | 86 |
| 25 nM | 1863 | 139559 | 75 |
| 12.5 nM | 696 | 65818 | 95 |
| 6.25 nM | 242 | 30546 | 126 |
| 3.125 nM | 168 | 15580 | 93 |
| 1.56 nM | 204 | 6630 | 33 |
| 0.78 nM | 156 | 3598 | 23 |
| 0.39 nM | 97 | 2170 | 22 |
| 0.19 nM | 325 | 1517 | 5 |
| 0.097 nM | 235 | 850 | 4 |
| 0.048 nM | 150 | 400 | 3 |

**Supporting Figure 4**. Serial dilution of productive ternary complex. (A) Luminescence signal plotted as a function of decreasing concentration of equimolar (1:1:1) mixture of $^{SUMO}$Sm-S1, $^{SUMO}$Lg-S2, and dNSP10 template (green bars). Background readings (RLU) collected from samples containing binary, equimolar mixture (1:1) of $^{SUMO}$Sm-S1, $^{SUMO}$Lg-S2 without template dNSP10 (black bars). (B) Observed values of signal, background, and signal/background ratio at each concentration tested. Note that with $^{SUMO}$Sm-S1, $^{SUMO}$Lg-S2 at 25 x $10^{-9}$ M, the *no template* samples (black bars) reproducibly gave NanoBiT bioluminescence that exceeded "blank" samples, containing buffer only (RLU ~100-200). This observation is consistent with a low level of spontaneous NanoBiT assembly. We took advantage of that template-independent NanoBiT signal as an internal positive control for the function of the two bioconjugates and the NanoBiT substrate in test samples.

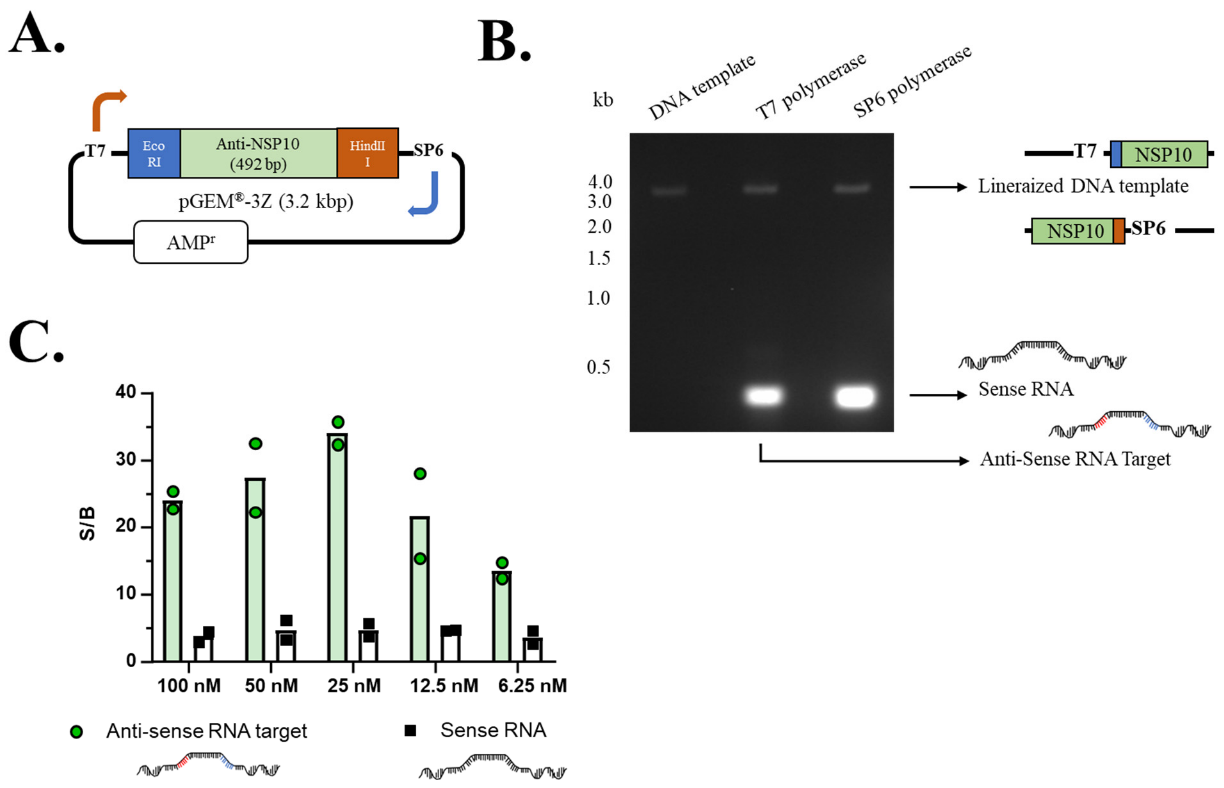

**Supporting Figure 5.** System for in vitro transcription of SARS-CoV-2 NSP10 sense and antisense RNA and "hook-effect" in sample assays. (A) Vector for in vitro transcription of NSP10. (B) In vitro transcribed sense and anti-sense NSP10 RNA separated by agarose gel electrophoresis and ethidium bromide staining. (C) Provisional evidence for bell shaped curve in fragment complementation assay. Luminescence signal from reconstituted NanoBiT in the presence of anti-sense target (green bars). Samples containing (+) strand NSP10 RNA show background signal.

### SUPPORTING TABLES

**Supporting Table 1**: Split enzyme-oligo complementation assay driven by hybridization

| Split Reporter | Oligo length in reporter conjugate | means of attachment | fragment-oligo conjugate concentration in assay | Template | signal/ background | mismatch selectivity | Hook Effect | Ref. |
|---|---|---|---|---|---|---|---|---|
| split Nluc | 20 | DHhC catalysis | LrgBiT: 25 nM SmBiT: 25 nM | ssDNA / RNA | 30 to 120-fold | requires > 2 bases change | yes | this work |
| split Nluc | 36 | ncCas9 | LrgBiT: 10 nM SmBiT:100 nM | dsDNA | 50-fold | not discussed | yes | van der Veer, Harmen J., et al. |
| split Nluc | 10 and 20 | Snap-tag | LrgBiT: 50 nM SmBiT: 200 nM | ssDNA | 175-fold | 50% decrease with 1-3 mismatches (using 10 mer) | not discussed | Zhou, Lanlan, et al. |
| Split mDHFR | 16 | maleimide coupling | 100 nM | ssDNA | 30-fold | 3-5 mismatches decrease rate 1.4 to 2.2-fold | yes | Sancho Oltra, Núria, Jeffrey Bos, and Gerard Roelfes. |

### SUPPORTING METHODS

#### 1. Materials

##### 1A. Chemicals

All chemicals were acquired from commercial suppliers and used directly: N,N-diisopropylethylamine, imidazole (Acros); N,N,N',N'-tetramethyl-O- (N-succinimidyl) uronium tetrafluoroborate, 1,7-diaminoheptane, ethylenediaminetetraacetic acid (Aldrich); Foscholine-12 (ANATRACE); GelRed (Biotium); Tris base (CORNING); HPLC grade water in 0.1% TFA, HPLC grade acetonitrile in 0.1% TFA (EMD Millipore Corp.); Ac-dC-CPG 500, dA-CPG-500, dA-CE phosphoramidite, Ac-dC-CE phosphoramidite, dmf-dG-CE phosphoramidite, dT-CE phosphoramidite, 5'-Carboxy-Modifier C10, Triethylammonium acetate (Glen Research); Isopropyl β-d-1-thiogalactopyranoside, tris (2-Carboxyethyl) phosphine hydrochloride (GOLDBIO); Methanol (MACRON); LB agar (miller); HPLC grade acetonitrile (Fisher Chemicals); Nano-Glo® Luciferase Assay System, Riboprobe® Combination Systems SP6/T7 (Promega); agar (Promega/Thermo); 23, 24-bisnor-5α-cholenic acid-3β-ol (Steraloids); Luria Bertani broth (Sigma); KCl, $MgCl_2$ (VWR)

##### 1B. Instruments for Synthetization, Purification and NanoBiT Detection

Incubator: Innova 42
Centrifuge: Multifuge X3 FR (Thermo)
Ni-NTA (Cytiva)
SDS-PAGE running apparatus: Mini-protein Tetra System (BIORAD)
Agarose gel running apparatus: Mini-sub cell GT (BIORAD)
Imager: Gel Doc EZ Imager (BIORAD)
HPLC: L-7400 (HITACHI)
C18 (RESTEK Viva C18 5 µm 250 x 4.6 mm)
C4 (Symmetry300™ C4 3.5 µm 4.6x150mm)
Corning® 96 Well Black Polystyrene Microplate (#3650)
Plate reader: Biotek Synergy H1
Solid phase synthesis: Expedite 8909 DNA/RNA synthesizer

##### 1C. Buffers

**Supporting Table 2:** Buffers and recipes

| Bacterial Cell Lysis Buffer | 2x Ni-NTA Binding Buffer | Ni-NTA Wash Buffer | Ni-NTA Elution Buffer |
|---|---|---|---|
| 0.05 M K2HPO4<br>0.4 M NaCl<br>0.1 M KCl<br>0.01 M imidazole<br>10% glycerol<br>0.5% Triton X-100<br>pH=7.4 | 1 M NaCl<br>0.04 M Na2HPO4<br>0.06 M Imidazole<br>20% glycerol<br>pH=7.4 | 0.5 M NaCl<br>0.02 M Na2HPO4<br>0.075 M Imidazole<br>10% glycerol<br>pH=7.4 | 0.5 M NaCl<br>0.02 M Na2HPO4<br>0.5 M Imidazole<br>10% glycerol<br>pH=7.4 |
| TE Buffer | 4x NanoBiT Assay Buffer | Agarose Gel Extraction Buffer | 5x BEN Buffer |
| 1 M Tris<br>0.5 M EDTA<br>pH=8.0 | 0.4 M KCl<br>0.004 M $MgCl_2$<br>0.04 M Tris<br>pH=8.0 | 0.02 M Tris<br>0.05 M NaCl<br>pH=7.4 | 0.1 M Bis-Tris<br>0.025 M EDTA<br>0.5 M NaCl<br>pH=7.4 |

#### 2. Methods:

##### 2A. Protein expression and purification

Two precursor proteins: SUMO-SmBiT-DHhC-His$_6$ and SUMO-LrgBiT-DHhC-His$_6$, were expressed by *E. coli* BL21(DE3) using the pET22b expression vector. In this plasmid, DHhC was subcloned using NcoI and HindIII; a synthetic fragment for SUMO-SmBiT or SUMO-LrgBiT was subcloned using the vector's NdeI and NcoI sites. *E. coli* expression cultures were grown in 50 mL sterilized LB broth plus carbenicillin (100 µg/mL, final) with 250 RPM shaking at 37 °C. When the OD600 reached 0.6-0.8, IPTG (0.5 mM, final) was added to induce protein expression and the temperature was reduced to 16 °C. Bacteria were harvested after 18-20 hours by centrifugation at 10,000 RPM for 10 min. The resulting pellet was resuspended in 3 mL of lysis buffer (+ lysozyme). After one -80 °C freeze/thaw cycles, DNase-I (3-5 µL) was added to the suspension followed by vortexing and sonication. Once the lysate appeared free flowing, the suspension was centrifuged at 10,000 RPM for 1 hour to pellet insoluble debris. The total soluble protein was then transferred to a clean 50 mL tube and combined with an equal volume of ice-cold 2x Ni-NTA binding buffer. The His-tagged protein was purified over a pre-equilibrated Ni-NTA spin column (Cytiva) essentially in accord with the manufacturer's instructions. Recipes for the buffers can be found in **Supporting Table 2**. Columns were washed 3x-5x with the Ni-NTA Wash Buffer, 500 µL each wash. The protein of interest was eluted using Ni-NTA elution buffer (350 µl)

##### 2B. Sterylamine Synthesis and Purification

Sterylamine (**III**) was prepared by amide coupling. To 4 mL anhydrous DMF at room temperature, we added 23, 24-bisnor-5α-cholenic acid-3β-ol (**I**), 34.7 mg (0.1 mmol) followed by N, N, N', N'-tetramethyl-O- (N-succinimidyl) uronium tetrafluoroborate (TSTU) 33.12 mg (0.1 mmol) and N,N-diisopropylethylamine (DIPEA) 87 µL (0.5 mmol). After 30 minutes of stirring, we added 1 mL (0.3 mmol) of 1,7-diaminoheptane (**II**) dropwise and continued mixing overnight at room temperature.

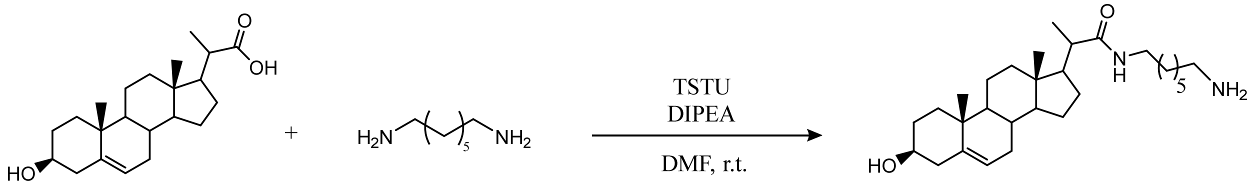

I II III

The reaction mixture was dried under vacuum and resuspended in pure methanol for reverse phase HPLC purification. To separate pure mono-sterylated diaminoheptane (**III**), we used a C18 column (RESTEK Viva C18 5 µm 250 x 4.6 mm) with a solvent gradient using HPLC grade water containing 0.1% trifluoroacetic acid (TFA) as buffer A; HPLC grade acetonitrile with 0.1% TFA as buffer B. Prior to sample injection, the column, injection loop, and injection needle were equilibrated with 100% buffer A. The gradient elution started with 100% buffer A and progressed to 100% buffer B over a period of 30 minutes. The detection wavelength was set at 210 nm. Sterylamine (**III**) eluted at 13 minutes.

##### NMR Characterization of Purified Sterylamine

Purified sterylamine (**III**) was redissolved in methanol- d4, 1H NMR and 13C NMR spectra was obtained with Bruker Avance III HD 400, proton chemical shifts are reported as δ in parts per million (ppm) relatives to methanol-d4

$^1$H NMR (400MHz, $CD_3OD$) δ 5.36 (dd, 1H), 3.41 (m, 1H), 3.18 (m, 1H), 3.13 (m, 1H), 2.92 (t, 2H), 1.40 (s, 6H), 1.16 (d, 4H), 1.05 (s, 3H), 0.76 (s, 3H). $^{13}$C NMR (400MHz, $CD_3OD$) δ 178.2, 140.9, 120.90, 71.0, 56.5, 52.6, 50.3, 43.7, 42.0, 41.6, 39.6, 39.3, 38.5, 37.1, 36.3, 31.9, 31.6, 30.9, 28.9, 28.3, 27.1, 27.0, 26.3, 26.0, 23.9, 20.7, 18.5, 16.5, 11.1

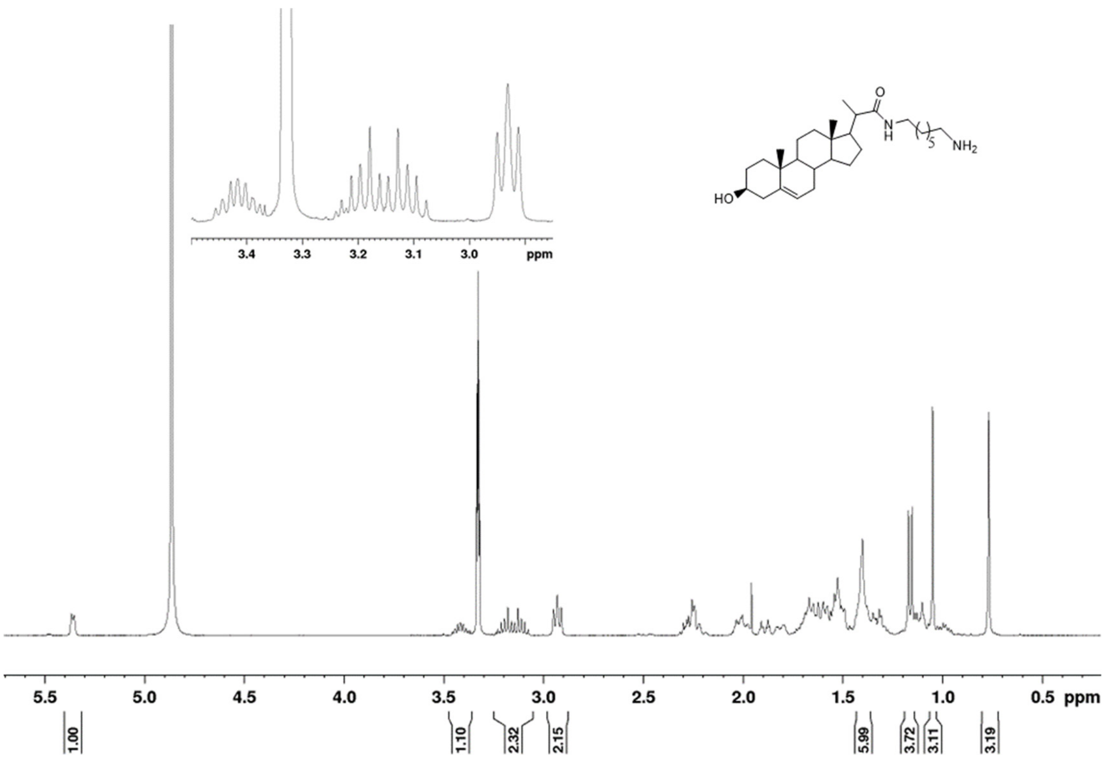

**Supporting Figure 6**. $^{1}$H NMR spectrum of sterylamine (**III**)

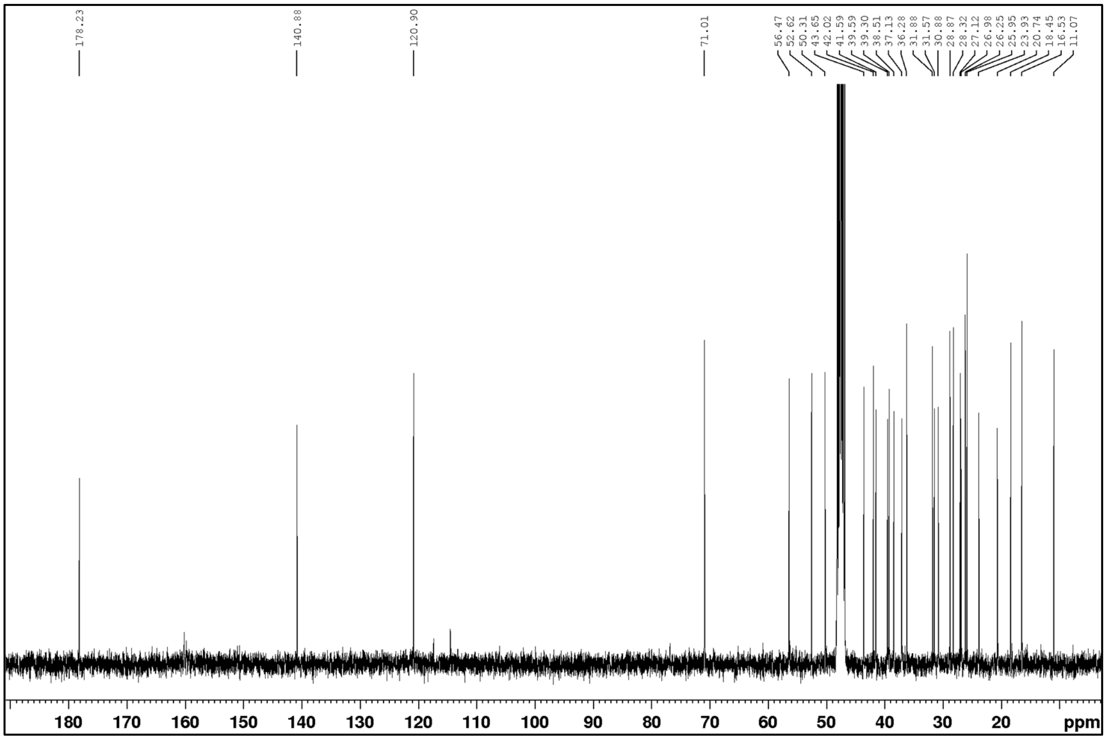

**Supporting Figure 7.** $^{13}$C NMR of the synthetic sterylamine (**III**)

**2C. Synthesis of Steramers and Purification:**
**Solid Phase Steramer Synthesis**

The sterylated-DNA oligomers (steramers) (**V**) were produced on a 1.0 µmole scale using standard DNA synthesis methods on an automated Expedite 8909 DNA/RNA synthesizer. Steramer 1 was synthesized using Ac-dC-CPG 500, while steramer 2 was synthesized using dA-CPG-500. Once the desired DNA sequence was synthesized, the 5'-end was linked with a 5'-carboxy modifier-C10 carrying a reactive NHS ester. After coupling with the 5'-carboxy modifier-C10, the detritylation step was performed. Before proceeding to the next coupling involving sterol amine, the solid support was washed with acetonitrile and dried using nitrogen purging for 5 minutes.

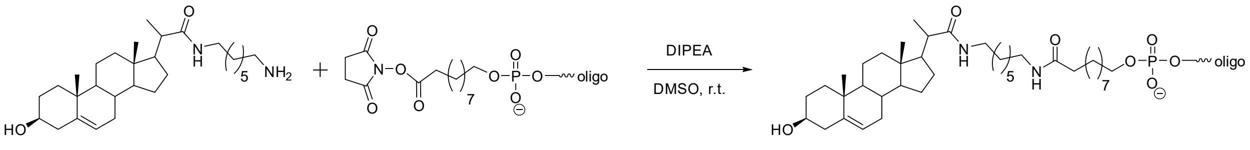

**III** **IV** **V**

For the manual coupling of sterylamine (**III**) with the oligo NHS ester (**IV**) on the CPG solid support, the procedure was carried out under anhydrous conditions. Anhydrous DMSO (0.05 M in **III**) containing 10% DIPEA was used as the solvent at room temperature. The coupling reaction took place for 2 hours using the push-pull syringe method. To prepare the coupling cocktail, sterylamine **III** (21 mg, 45 µmole) was dissolved in 810 µL of dry DMSO. Then, 90 µL of dry DIPEA was added to the solution, resulting in a total volume of 900 µL for the coupling cocktail. For each 1 µmole synthesis, 200 µL of the coupling cocktail was used. Following the coupling step, the column was thoroughly washed with 5 × 1 mL of acetonitrile and subsequently dried.

The steramers were detached from the solid support by treating them with 40% methyl amine in water (2 × 0.8 mL) at room temperature for 2 hours. After the cleavage, the resulting solution was subjected to lyophilization, which yielded the crude steramers.

**Steramer Purification by C4 HPLC:**

HPLC grade 0.1 M TEAA in water was used as buffer A; HPLC grade 0.1 M TEAA in 50% acetonitrile was used as buffer B. A gradient elution was applied, starting with 20% buffer B and progressing to 100% buffer B over a period of 30 minutes. 100% Buffer B remained for 5 min, then decreased back to 20% in 5 min. The detection wavelength was set at 260 nm. Steramer fractions were collected and dried under nitrogen, then resuspended with TE buffer. The purified sample was reinjected to confirm the purity. To confirm the identity of the steramers, we used MALDI-ToF analysis (**Supporting Table 3 and Supporting Figure 8 and 9**).

**Supporting Table 3**: List of steramers synthesized by solid phase synthesis.

| Name | Sequence | Calculated Mass | Mass found |
|---|---|---|---|
| Steramer 1 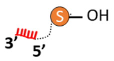 | sterol-5’-TTATGGCTGTAGTTGTGATC-3’ | 6860 | 6860 |
| Steramer 2 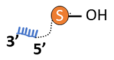 | sterol-5’-CGTCTGCGGTATGTGGAA-3’ | 6262 | 6263 |

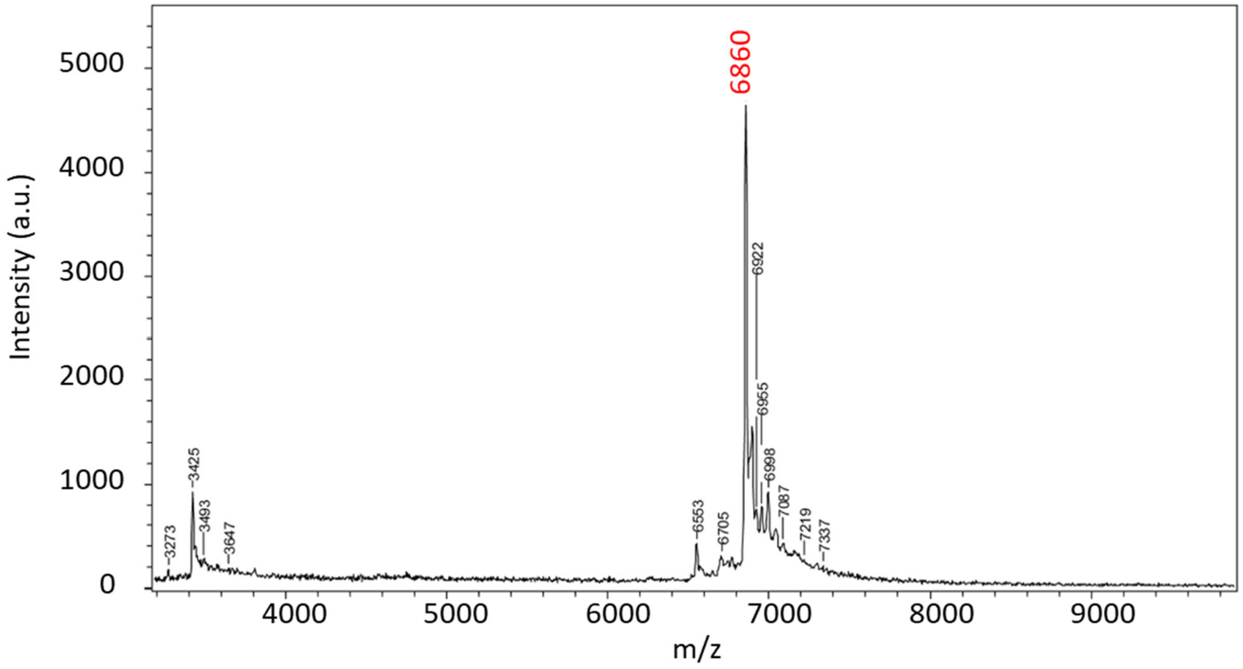

**Supporting Figure 8.** MALDI-ToF analysis of streamer 1

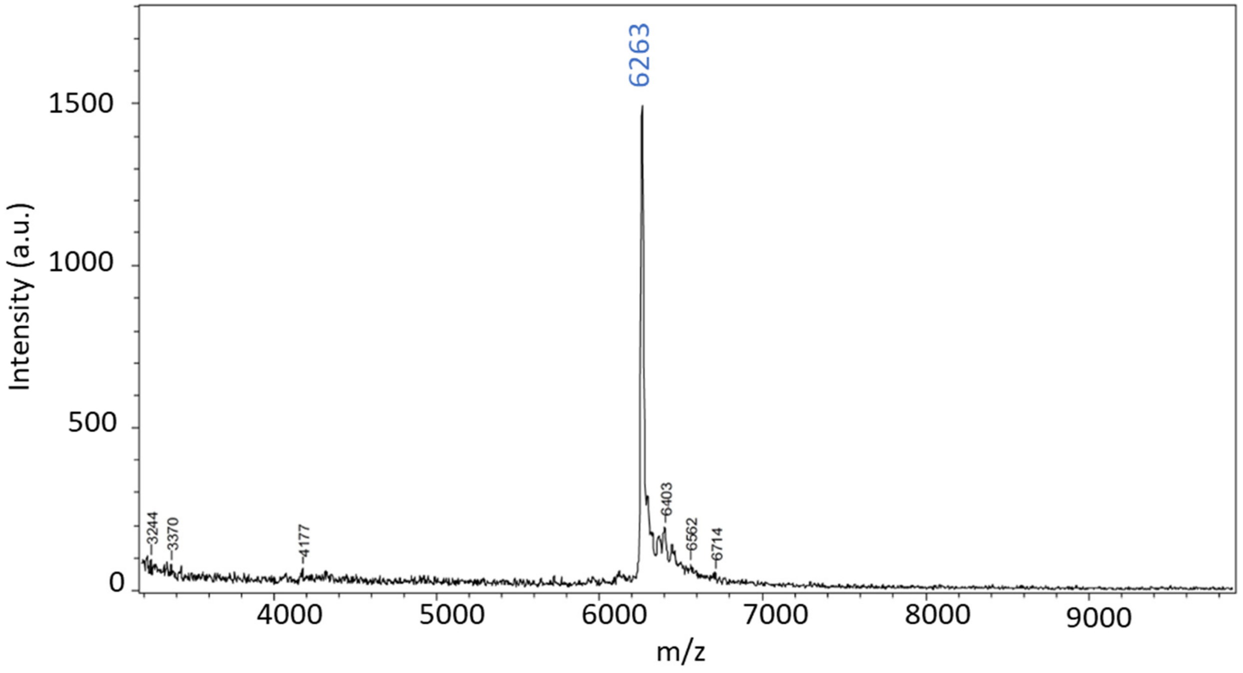

**Supporting Figure 9.** MALDI-ToF analysis of streamer 2

**2D. Conjugate Preparation and Purification:**
**Conjugation Reaction**

To obtain the protein-nucleic acid conjugates used here, bench top reactions were performed in DHhC activity buffer (Tris Buffer pH 7.4 (0.02 M, final), Fos-choline 12 (0.0015 M, final), Tris (2-Carboxyethyl) phosphine Hydrochloride (TCEP) pH 7.4 (0.001 M, final). Precursor proteins (SUMO-SmBiT-DHhC, SUMO-LrgBiT-DHhC) were added to

$1.5 \times 10^{-6}$ M, final followed by streamer 1 or 2 ($5 \times 10^{-5}$ M, final). The reaction mixture was incubated at room temperature, typically overnight.

**Agarose gel Extraction purification of NanoBit Steramer sensors:**

To the reaction mixture, an appropriate amount of gel loading dye, purple (6X) was added. A 2% agarose with GelRed nucleic acid stain was used to separate the conjugate from precursor protein and unreacted steramer. A representative gel is shown (**Supporting Figure 10**). Gel slices containing the conjugates were removed and extracted with 20 mM Tris pH 7.4, 50 mM NaCl buffer overnight. Any remaining agarose was removed by centrifugation.

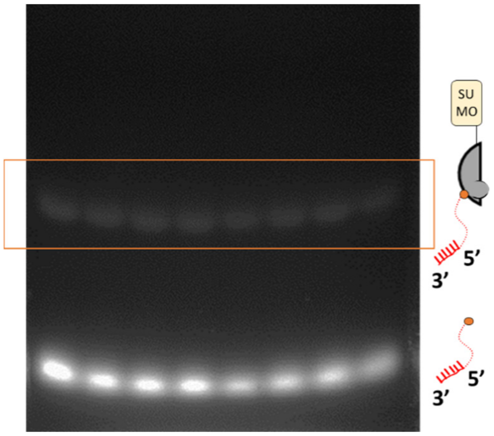

**Supporting Figure 10:** Agarose gel separation of the indicated conjugate after staining with Gel-Red. The lower bands represent excess steramers.

**2E. FRET Analysis of steramer substrate activity with DHhC**

Reactions were carried out in 96-well Corning black plates. Prior to adding substrate, the initial reaction volume of 96 µL contained Bis-Tris buffer (0.05 M, final, pH 7.1), ethylenediaminetetraacetic acid (EDTA, $5\times10^{-4}$ M, final), NaCl (0.1 M, final), Fos-choline 12 (0.0015 M, final), Tris (2-Carboxyethyl) phosphine Hydrochloride (TCEP) pH 7.4 (0.001 M, final) and C-H-Y ($2 \times 10^{-7}$ M, final), as described previously (Owens et. al 2005). After temperature equilibration at 30 °C for 10 minutes, 4 µL of cholesterol, or steramer was added to initiate reaction. FRET ratio of 540 nm/460 nm was recorded using a Biotek Synergy H1 plate reader. Kinetic traces of C-H-Y FRET loss were fit to a first order decay using excel solver to calculate $k_{obs}$.

$$\mathrm{FRET} = \mathrm{A} * \mathrm{e}^{-k_{\mathrm{obs}}\mathrm{t}} + \mathrm{C}$$

The substrate binding affinity for DHhC was determined by Michaelis-Menten plots. With the same concentration of buffers and C-H-Y described above, substrates were assayed over a dilution series from final $5 \times 10^{-5}$ M to $2 \times 10^{-6}$ M. The initial velocity of FRET loss at each concentration was determined by the slope of the linear portion of the (FRET vs time) plots. Data were used to find $V_{max}$ and $K_M$ values in the following equation using excel solver.

$$V_{obs} = \frac{V_{max} * [S]}{K_M + [S]}$$

**Supporting Table 4**: Amino acid and DNA sequences and annotation of bioconjugates and nucleic acid targets

| | |
|---|---|
| 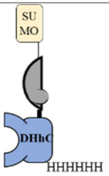<br>SUMO-SmBiT-<br>DHhC | MGSDSEVNQEAKPEVKPEVKPETHINLKVSDGSSEIFFKIKKTTPLRRLMEAFAKRQ<br>GKEMDSLRFLYDGIRIQADQTPEDLDMEDNDIIEAHREQIGGGSGMVTGYRLFEEIL<br>GSSGGGGSGGGGHGCFTPESTALLESGVRKPLGELSIGDRVLSMTANGQAVYSEVIL<br>FMDRNLEQMQNFVQLHTDGGAVLTVTPAHLVSVWQPESQKLTFVFADRIEEKNQV<br>LVRDVETGELRPQRVVKVGSVRSKGVVAPLTREGTIVVNSVAASCYAVINSQSLAH<br>WGLAPMRLLSTLEAWLPAKEQLHSSPKVVSSAQQQNGIHWYANALYKVKDYVLP<br>QSWRHDGSGHHHHHHH |
| 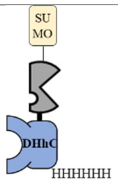<br>SUMO-LrgBiT-<br>DHhC | MGSDSEVNQEAKPEVKPEVKPETHINLKVSDGSSEIFFKIKKTTPLRRLMEAFAKRQ<br>GKEMDSLRFLYDGIRIQADQTPEDLDMEDNDIIEAHREQIGGGSGGGSGGSGVFTLE<br>DFVGDWEQTAAYNLDQVLEQGGVSSLLQNLAVSVTPIQRIVRSGENALKIDIHVIIP<br>YEGLSADQMAQIEEVFKVVYPVDDHHFKVILPYGTLVIDGVTPNMLNYFGRPYEGI<br>AVFDGKKITVTGTLWNGNKIIDERLITPDGSMLFRVTINSGSSGGGGSGGGGHGCFT<br>PESTALLESGVRKPLGELSIGDRVLSMTANGQAVYSEVILFMDRNLEQMQNFVQLH<br>TDGGAVLTVTPAHLVSVWQPESQKLTFVFADRIEEKNQVLVRDVETGELRPQRVVK<br>VGSVRSKGVVAPLTREGTIVVNSVAASCYAVINSQSLAHWGLAPMRLLSTLEAWLP<br>AKEQLHSSPKVVSSAQQQNGIHWYANALYKVKDYVLPQSWRHDGSGHHHHHHH |
| SmBiT Conjugate<br>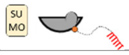 | SUMO-SmBiT-sterol linker-5’-TTATGGCTGTAGTTGTGATC-3’ |
| LrgBiT Conjugate<br>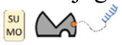 | SUMO-LrgBiT-sterol linker-5’-CGTCTGCGGTATGTGGAA-3’ |
| d-NSP10 | 5’-TTGATCACAACTACAGCCATAACCTTTCCACATACCGCAGACGGT-3’ |
| d-NSP10 $(TT)_1$ | 5’-TTGATCACAACTACAGCCATAACTTCTTTCCACATACCGCAGACGGT-3’ |
| d-NSP10 $(TT)_2$ | 5’-TTGATCACAACTACAGCCATAACTTTTCTTTCCACATACCGCAGACGGT-3’ |
| d-NSP10 $(TT)_3$ | 5’-TTGATCACAACTACAGCCATAACTTTTTTCTTTCCACATACCGCAGACGGT-3’ |
| d-NSP10 $(TT)_4$ | 5’-TTGATCACAACTACAGCCATAACTTTTTTTTCTTTCCACATACCGCAGACGGT-<br>3’ |
| d-NSP10 (A to C) | 5’-TTGATCACAACTACAGCCATAACCTTTCCACATCCCGCAGACGGT-3’ |
| d-NSP10 (C to A) | 5’-TTGATCACAACTACAGCCATAACCTTTCCACATAACGCAGACGGT-3’ |
| d-NSP10 (AC swap) | 5’-TTGATCACAACTACAGCCATAACCTTTCCACATCACGCAGACGGT-3’ |
| SP6 Polymerase RNA product (sense) | 5’-<br>UUAUGAGUUCGAAAUUAACACAAUUCUACAACACAUGUGUGUGACCAUGACC<br>AGUCCGUUAUUGUCAAUGUGGCCUUCGGUUAUACCUAGUUCUUAGGAAACCA<br>CCACGUAGCACAACAGACAUGACGGCAACGGUGUAUCUAGUAGGUUUAGGAU<br>UUCCUAAAACACUGAAUUUUCCAUUCAUACAUGUUUAUGGAUGUUGAACACG<br>AUUACUGGGACACCCAAAAUGUGAAUUUUUGUGUCAGACAUGGCAGACGCCA<br>UACACCUUUCCAAUACCGACAUCAACACUAGUUGAGGCGCUUGGGUACGAAG<br>UCAGUCGACUACGUGUUAGCAAAAAUUUGCCCAAACGCCACAUUCACGUCGG<br>GCAGAAUGUGGCACGCCGUGUCCGUGAUCAUGACUACAGCAUAUGUCCCGAA<br>AACUGUAGAUGUUACUAUUUCAUCGACCAAAACGAUUUAAGGAUUUUUGAUU<br>AACAACAGCGAAGGUUCUUUUCCUGCUUAAG-3’ |
| T7 Polymerase RNA product (anti-sense) | 5’-<br>CGAAUUCGUCCUUUUCUUGGAAGCGACAACAAUUAGUUUUUAGGAAUUUAGC<br>AAAACCAGCUACUUUAUCAUUGUAGAUGUCAAAAGCCCUGUAUACGACAUCA<br>GUACUAGUGCCUGUGCCGCACGGUGUAAGACGGGCUGCACUUACACCGCAAA<br>CCCGUUUAAAAACGAUUGUGCAUCAGCUGACUGAAGCAUGGGUUCGCGGAGU<br>UGAUCACAACUACAGCCAUAACCUUUCCACAUACCGCAGACGGUACAGACUG<br>UGUUUUUAAGUGUAAAACCCACAGGGUCAUUAGCACAAGUUGUAGGUAUUUG<br>UACAUACUUACCUUUUAAGUCACAAAAUCCUUUAGGAUUUGGAUGAUCUAUG<br>UGGCAACGGCAGUACAGACAACACGAUGCACCACCAAAGGAUUCUUGAUCCA |

| | |
|---|---|
| | UAUUGGCUUCCGGUGUAACUGUUAUUGCCUGACCAGUACCAGUGUGUGUACA CAACAUCUUAACACAAUUAAAGCU-3' |

**2F. Bioluminescence Measurement and Data Analysis:**
For protein complementation reactions, NanoBiT conjugates and nucleic acid targets were mixed in hybridization buffer (75 µL total), incubated for 10-30 minutes at 25 °C, then assayed for NanoBiT enzymatic activity by adding 25 µL of Nano-Glo® Luciferase Assay System substrate. Bioluminescence was measured using Biotek Synergy H1.

**Dose Response fitting to Bell-Shape Curve**

For hook effect experiments, the concentration of both NanoBiT conjugates was $25 \times 10^{-9}$ M, while the nucleic acid template varied from $3.2 \times 10^{-6}$ M to $1.9 \times 10^{-10}$ M. The resulting concentration response plots showed a bell-shape curve that could be described by the equation:

$$Y = Dip + \frac{(Plateau_1 - Dip)}{1 + 10^{(\,Log\ EC50_1 - X) * Hill\ slope_1}} + \frac{(Plateau_2 - Dip)}{1 + 10^{(X - \,Log\ EC50_2) * Hill\ slope_2}}$$

**2G. RNA Preparation and Purification**
The NSP10 gene was inserted in the 3' to 5' direction into pGEM®-3Z vector (Promega) using EcoRI and HindIII. RNA was transcribed by using Riboprobe® Combination Systems SP6/T7 (Promega), 20 µL total reaction mixture with 0.2 µg of linearized DNA template. Reactions were incubated at 37 °C for 2 hours and followed with DNase treatment at 37 °C for 30 min. The RNA was then purified using Monarch® Kits for RNA Cleanup (NEB) and eluted with 50 µL of RNase -free buffer. A typical yield was 10 µg of RNA.
